# Folic acid prevents *C. elegans* folate deficiency indirectly via bacterial uptake of a breakdown product: a route that can also increase bacterial toxicity and ageing

**DOI:** 10.1101/230227

**Authors:** Claire Maynard, Ian Cummins, Jacalyn Green, David Weinkove

## Abstract

Supplementation with the synthetic oxidised folate, folic acid is used to prevent neural tube defects and other symptoms of folate deficiency. However, several unanswered questions remain over folic acid efficacy, safety and interactions with gut microbes. Prevention of a development defect caused by folate deficiency in the nematode worm *Caenorhabditis elegans* requires > 10 fold higher concentrations of folic acid compared to folinic acid, a reduced folate. Here we show that the major route for folic acid to restore normal development is indirect via the *Escherichia coli* used to feed *C. elegans.* This route occurs mainly via the *E. coli* transporter AbgT, which takes up the folic acid breakdown product para-aminobenzoate-glutamate (PABA-glu). We found that folic acid preparations, including a commercial supplement, contain 0.3- 4.0 % of this breakdown product. Previously, we have shown that inhibiting bacterial folate synthesis increases *C. elegans* lifespan by removing a life-shortening bacterial activity. Here, we show that folic acid restores bacterial folate synthesis and reverses this lifespan increase. It is still to be determined whether this bacterial route increases host folate levels in humans and if there are situations where increased bacterial folate synthesis has negative health complications.

## INTRODUCTION

The folate cycle is a set of essential biosynthetic reactions known as one carbon metabolism (Ducker and Rabinowitz, 2017). Folates are a family of molecules with a central aromatic core derived from para-amino benzoic acid (PABA), a pterin ring that can be modified and chain of one or more glutamates (Green and Matthews, 2007). At each step of the folate cycle, an enzyme mediates a modification of the pterin ring of the bound folate that allows the transfer of a chemical group containing one carbon atom (methyl, formyl etc.) to or from the compound being synthesised (Ducker and Rabinowitz, 2017). Because of this cofactor role for folate molecules, there are recycled and only required in very small amounts. Animals cannot synthesise folates and must acquire them from their diet or gut microbes. A symptom of human folate deficiency is neural tube defects during embryonic development (Ducker and Rabinowitz, 2017). It has been found the rate of these defects can be lowered by preconception supplementation with folic acid, an oxidised form of folate not found in nature. In many countries including the US and Canada, there is mandatory fortification of flour with folic acid and this intervention has successfully decreased the incidence of birth defects (Bailey et al., 2015). There are some concerns that folic acid supplementation may have adverse effects (Kim, 2007; Marean et al., 2011; Pickell et al., 2011; Smith et al., 2008), and there are many unknowns about the efficacy of uptake and biological utilisation of the supplement (Gregory et al., 2005). However, recent reviews of the evidence by experts acting for government public health bodies have concluded that the risks are minimal and have recommended either the supplementation of flour or other food products as a beneficial intervention (Bailey et al., 2015; Boyles et al., 2016; Public_Health_England, 2017). These reviews do not mention the potential role of gut microbes in considering the mechanisms and safety of folic acid supplementation.

Research from our group and others has shown that inhibiting folate synthesis in *Escherichia coli*, either by treatment with the drug sulfamethoxazole (SMX) or mutation of the PABA synthesis pathway (e. g. a *pabA* or *pabB* mutant), extends the lifespan of the nematode *Caenorhabditis elegans* that feeds on it (Han et al., 2017; Virk et al., 2012; Virk et al., 2016). While these interventions decrease the folate levels in both *E. coli* and *C. elegans,* there remains sufficient folate available to support normal growth of both (Virk et al., 2012). Numerous lines of evidence suggest that lifespan is increased because inhibiting folate synthesis prevents a life-shortening activity of the bacteria rather than because animal folate levels are decreased (Virk et al., 2016). A mutant in the *C. elegans* homologue of the human GCPII enzyme (which cleaves glutamates from polyglutamated folates (Halsted et al., 1998)) shows delayed development and infertility on low folate *E. coli* (Virk et al., 2016). This defect is due to folate deficiency because it can be prevented by adding 1-10 μM folinic acid, a reduced folate. Prevention with folic acid requires much higher concentrations (100 μM) (Virk et al., 2016). This result may reflect the fact that the only characterised *C. elegans* folate transporter is FOLT-1, a reduced folate carrier that shows specificity for folinic acid (Balamurugan et al., 2007). However, we also discovered that at high concentrations folic acid can partially reverse the lifespan increase caused by inhibiting *E. coli* folate synthesis. This finding suggests that *E. coli* can take up folic acid (Virk et al., 2012). *E. coli* does not possess a folate transporter (Nickerson and Webb, 1956; Webb, 1955) but can take up the folic acid breakdown product PABA-glu through the transporter AbgT and catabolise it to PABA (Carter et al., 2007). PABA can also diffuse across *E. coli* membranes. PABA from either source can be used to synthesise folate.

In this study, we show that the prevention of delays in *C. elegans* development by folic acid depends on the *E. coli* AbgT transporter, demonstrating that bacterial uptake is the major route of folic acid supplementation. Consistent with this finding, we show that folic acid increases *E. coli* folate levels through the AbgT transporter and we find PABA-glu present in folic acid preparations. We also show that uptake of PABA-glu by AbgT can reverse the lifespan increase caused by inhibiting *E. coli* folate synthesis. Thus, this pathway can have both positive effects on development and negative consequences for ageing.

## RESULTS

### The major route of folic acid prevention of a *C. elegans* developmental defect requires the *E. coli* AbgT transporter

In order to test whether folic acid requires *E. coli* to prevent developmental defects in our *C. elegans* folate deficiency model, we tested whether the *E. coli abgT* genotype influenced the outcome of supplementation. We first established that the *C. elegans gcp-2.1(ok1004)* mutant grown on the *E. coli pabA* mutant on defined media (DM) has the same growth defect as these worms grown on wild type *E. coli* treated with 128 μg/ml SMX (Figure 1a, Virk *et al.* 2016). Addition of 2.5 μM folinic acid or higher prevented this growth defect regardless of the *abgT* genotype (Figure 1b). In contrast, folic acid was found to increase *gcp-2.1* mutant body length on the *pabA* mutant in a dose-dependent manner, achieving full growth only at 200 μM folic acid (Figure 1c). On the *E. coli abgTpabA* double mutant, a much weaker response to folic acid was observed, whereas overexpression of *abgT* in the *E. coli pabA* mutant resulted in normal *C. elegans* growth at lower concentrations of folic acid (Figure 1c). Analysing the experiment by two-way ANOVA, we find that there is a significant interaction effect of *abgT* genotype (F=102.67, p<0.0001) and folic acid concentration (F=123.55, p<0.0001) on *C. elegans gcp-2.1* body length. Mutation of *abgT* alone did not influence growth of the *gcp-2.1* mutant (Supplementary Figure 1a). These results are consistent with folinic acid being taken up directly by the worm (Balamurugan et al., 2007), and that the major route of folic acid uptake requires *E. coli* and the *E. coli* AbgT transporter.

**Figure 1.**
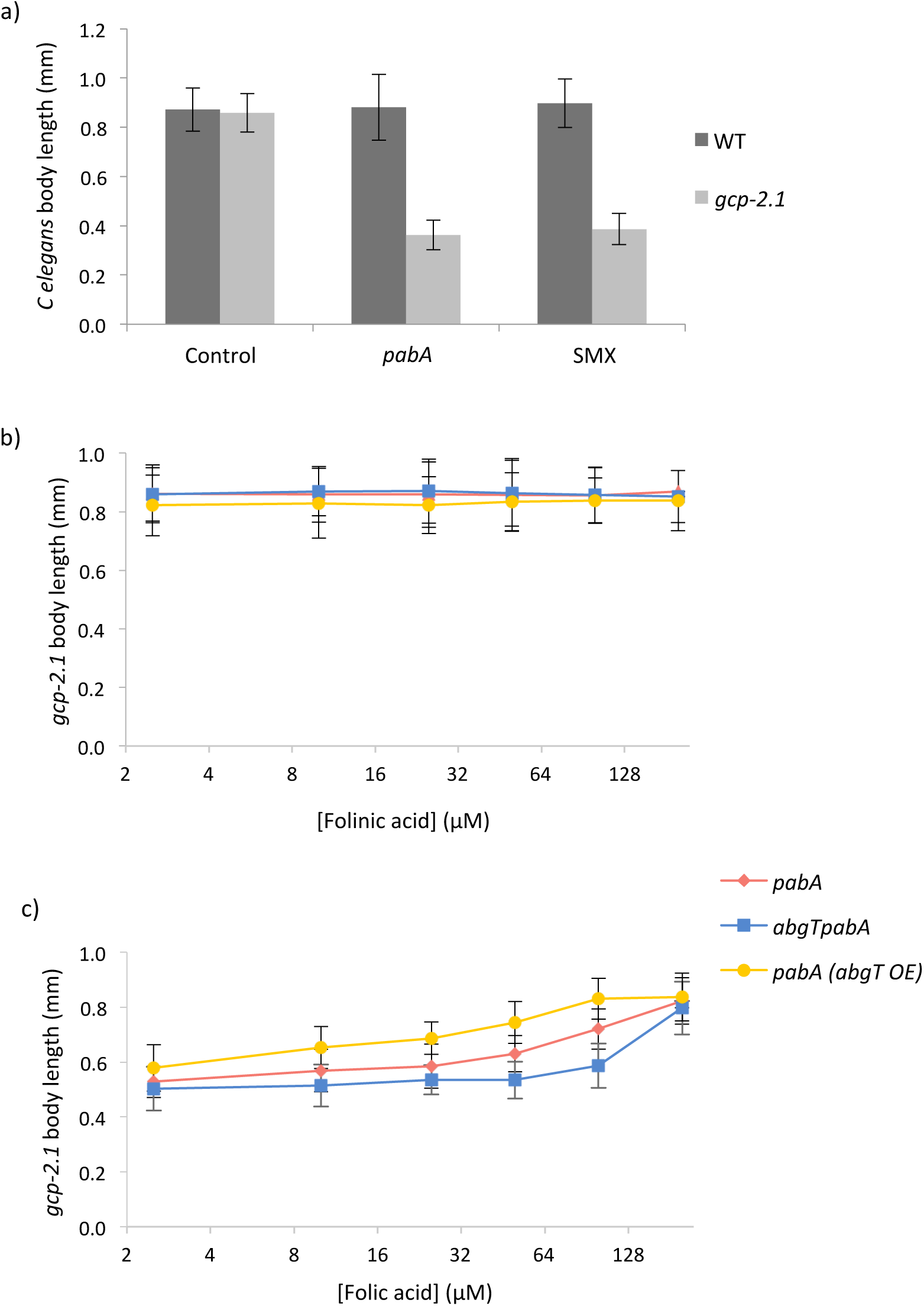
Folic acid prevents developmental delay of a *C. elegans* folate deficiency model via an *E. coli abgT*-dependent route. a) body length of wild-type and *gcp-2.1* mutant *C. elegans* at L4 stage raised on DM agar plates seeded with WT *E. coli* (control), *pabA* mutant or WT *E. coli* treated with 128 μg/ml SMX. b) body length of *gcp-2.1* mutant C. *elegans* at L4 stage raised on plates seeded with *pabA* mutant, *abgTpabA* double mutant or *pabA* mutant over-expressing *abgT* with increasing concentrations of folinic acid and c) folic acid: by two-way ANOVA analyses, we find that there is a significant interaction effect of strain type (F=102.67, p<0.0001) and folic acid concentration (F=123.55, p<0.0001) on C. *elegans gcp-2.1* body length. Over-expression is conferred by transformation with a high copy number plasmid, pJ128. *pabA* and *abgTpabA* strain are transformed with the empty vector, pUC19. Error bars represent standard deviation of C. *elegans* body length; n ≥40.

### Folic acid supports growth *of E. coli pabA* mutants via *abgT*-dependent uptake of PABA-glu

The dependence on *E. coli abgT* for folic acid to rescue the *gcp-2.1* developmental delay strongly suggests that PABA-glu is available to *E. coli* following folic acid supplementation. To test the relative ability of *E. coli* to take up PABA-glu and folic acid, we added these compounds to *pabA, abgT pabA* and *pabA (abgT OE) E. coli* and assessed growth on DM agar plates in PABA-free conditions under which the *pabA* mutant showed slower growth than wild type. We found that growth in the presence of folic acid and PABA-glu depended on *abgT* expression; 10 μM folic acid rescued growth of the *pabA* mutant, whereas 100 μM was needed to achieve an equivalent rescue in the *abgT pabA* double mutant (Figure 2a). In the presence of 10 μM folic acid, growth of the *pabA* strain over-expressing AbgT was greater than of the *pabA* mutant alone or the wild type. Supplementation by PABA-glu had a similar effect to folic acid but at a 10-fold lower concentration (Figure 2a). PABA, which can diffuse across biological membranes (Tran and Nichols, 1991), rescued bacterial growth at nanomolar concentrations independently of *abgT* expression (Figure 2a). Overall, rescue of bacterial growth by folic acid can be largely explained by PABA-glu uptake by AbgT while low concentrations of PABA present in folic acid preparations may explain the ability of 100 μM folic acid to rescue growth of the *E. coli abgT pabA* double mutant.

**Figure 2.**
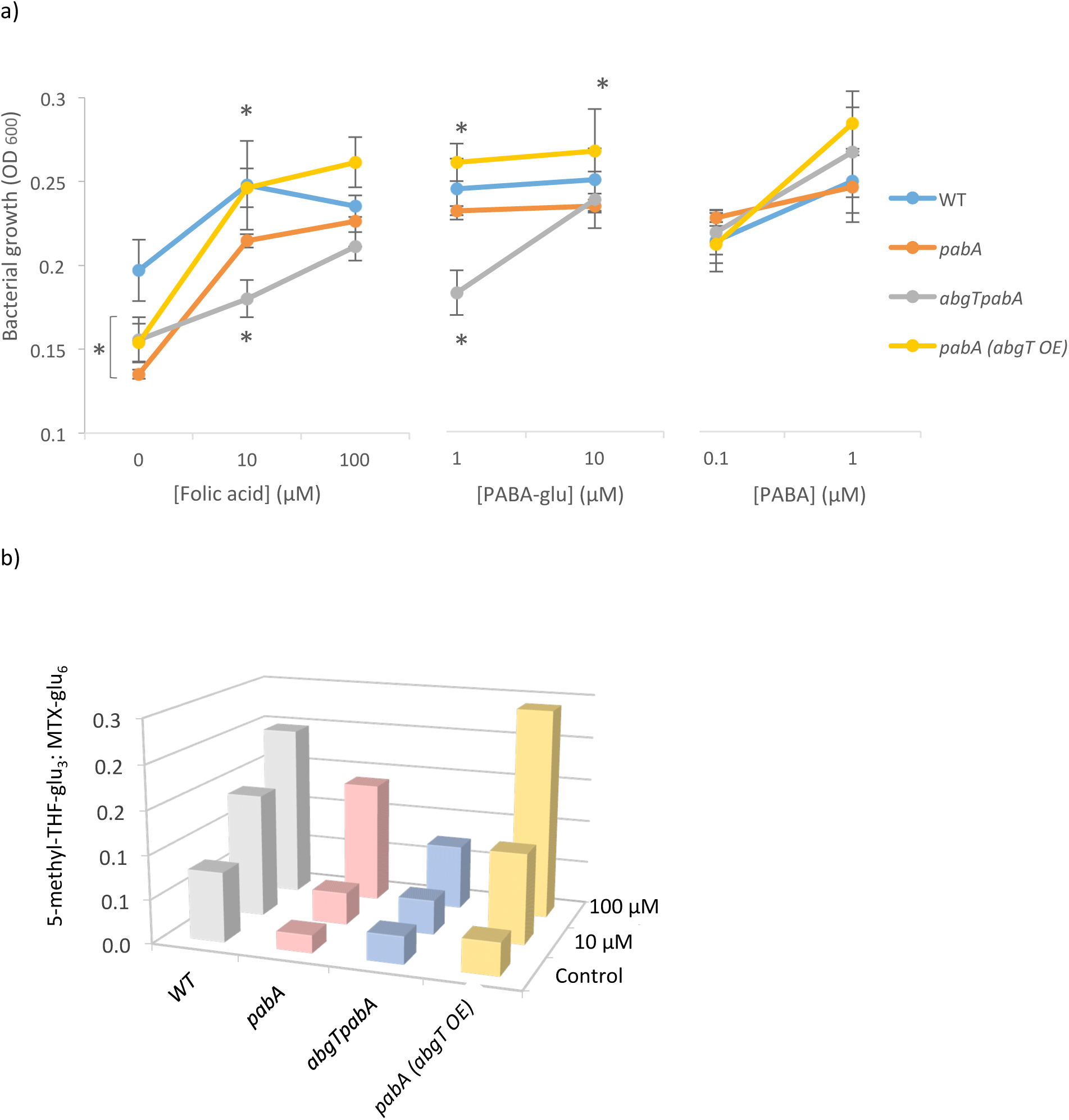
Folic acid breakdown supports *E. coli* growth and increases folate levels via uptake of PABA-glu by AbgT. a)growth of *E. coli* WT, *pabA, abgTpabA* and *pabA* over-expressing *abgT (abgT* OE) on agar plates supplemented with folic acid PABA-glu and PABA as measured by OD_600_ after 4 days growth at 25°C (see methods). Each data point is the average of 8 plates. Error bars represent standard deviation. Asterisks denote the test statistic from Student's *t* test comparison of means, where * = P<0.05 compared to WT growth on the same condition b) 5-methyl THF-glu3 levels in extracts of *E. coli WT, pabA, abgTpabA, pabA* (*abgT* OE) mutants supplemented with 10 μM and 100 μM folic acid. Folate counts from the LC-MS/MS are normalised by dividing by counts of an internal MTX-glu_6_ spike. Extracts were made after 4 days of bacterial growth at 25°C on solid agar plates. See Supplementary Figure 3 for full folate analysis.

### Folic acid increases *E. coli* folate levels in an AbgT-dependent mechanism

It order to verify that *E. coli* growth following folic acid supplementation is attributable to restored bacterial folate synthesis, we used LC-MS/MS to detect *E. coli* folate levels under the conditions used in the above experiment. Levels of the most detectable and thus likely the most abundant THF species, 5-methyl THF-glu_3_, are presented in Figure 2b. In the absence of folic acid, folate levels in the *pabA* mutant strains were significantly lower than in WT extracts (Figure 2b, Supplementary Figure 2). Addition of folic acid increased folate levels, where the scale of increase was dependent on *abgT* expression; with 100 μM folic acid, folate species were highest in *pabA* (*abgT* OE) followed by WT, *pabA,* and finally lowest in the *abgT pabA* double mutant. Other THF species followed the same trend as 5-methyl THF-glu_3_ and are presented in Supplementary Figure 2. In summary, folic acid supplementation increases *E. coli* folate levels in an *abgT*-dependent mechanism.

### Folic acid preparations contain PABA-glu and PABA

Together, the data presented here indicate that the main route of *C. elegans* folic acid supplementation is indirect via *E. coli* uptake of PABA-glu and PABA. We used LC-MS/MS to test for the presence of these breakdown products in folic acid preparations from Schircks (used in all other experiments in this study), Sigma Aldrich and Boots, a UK retailer. We also tested for further folic acid breakdown under the experimental conditions used here by analysing extracts from agar media supplemented with Schircks folic acid and incubated at 25 °C for 4 days. We detected PABA-glu in all three folic acid sources at between 0.3% (Schircks) and 4% (Boots, Figure 3). Under the conditions used for *C. elegans* experiments PABA-glu increased to 1.18%, suggesting further breakdown. PABA was found at between 0.01% (Schircks) and 0.06% (Boots, Figure 3).

**Figure 3.**
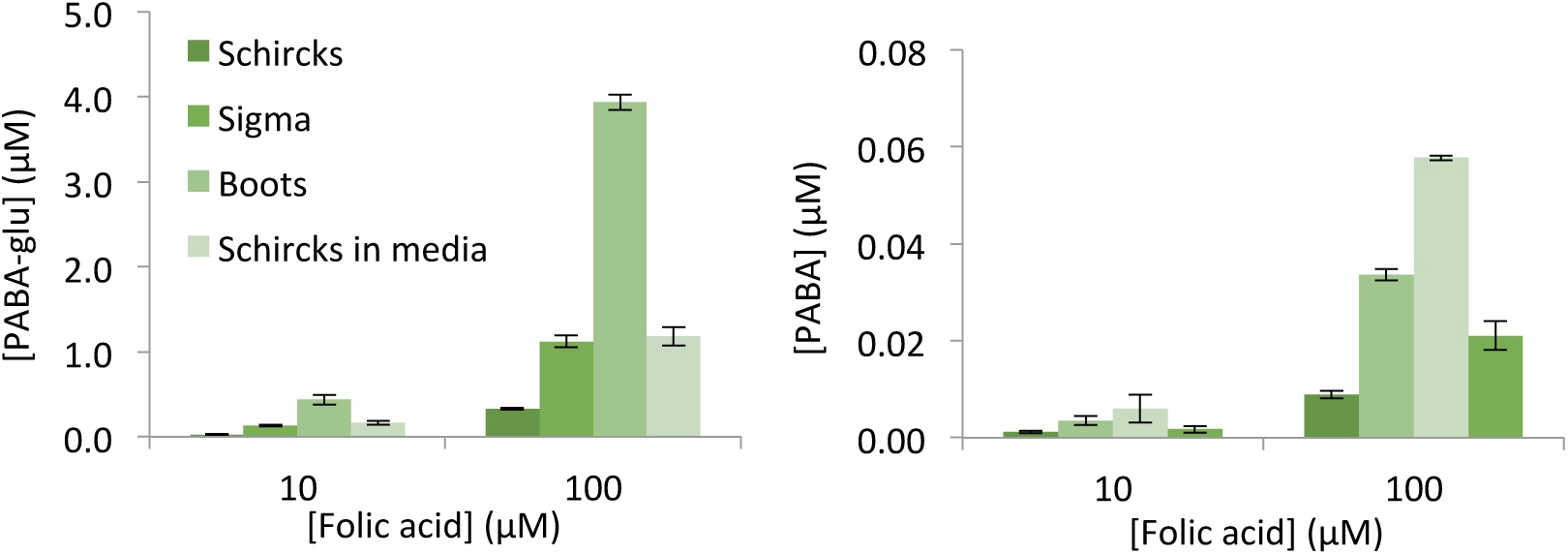
Folate preparations contain PABA-glu and PABA. Concentrations of PABA-glu and PABA as determined by LC-MS/MS in 10 and 100 μM folic acid preparations. Samples were folic acid from Schircks, Sigma, Boots and Schircks folic acid after addition to the agar media and incubation for 4 days at 25 °C. Error bars represent standard deviation over triplicate independent preparations.

### Folic acid shortens *C. elegans* lifespan via AbgT-dependent uptake of PABA-glu during adulthood

Inhibiting bacterial folate synthesis, without affecting bacterial growth, is known to increase *C. elegans* lifespan (Virk et al., 2012; Virk et al., 2016). It was therefore hypothesized that factors that increase bacterial folate synthesis, such as folic acid (as shown here), may shorten *C. elegans* lifespan. Consistent with our previous findings (Virk et al., 2016), we find that *C. elegans* maintained on any *E. coli pabA* mutant are long-lived compared to *C. elegans* fed wild type *E. coli* (Figure 4, Supplementary Table 1), whereas the *abgT* mutation alone had no impact on *C. elegans* lifespan (P=0.4312, Supplementary Figure 1b). 10 μM folic acid was found to decrease *C. elegans* lifespan on *pabA* by 9.4% (P=0.0052), and decrease lifespan on the *pabA* mutant over-expressing *abgT* further (by 16.3%, P<0.0001, Figure 4a). In contrast, 10 μM folic acid had no effect on lifespan on the *abgTpabA* double mutant (P=0.1901, Figure 4a). 100 μM folic acid decreased lifespan on *pabA E. coli* by 23.9% (P<0.0001), whereas this concentration only shortened lifespan on the *abgT pabA* double mutant by 4.7% (P=0.0467, Figure 3a). Lifespans on media supplemented with PABA-glu showed an abgT-dependent response similar to that observed with folic acid supplementation, but at a 10-fold lower concentration (Figure 4b). In contrast, PABA supplementation shortened *C. elegans* lifespan in all cases independently of *abgT* expression, consistent with its ability to diffuse across biological membranes (Figure 4c). Folic acid had no effect on the lifespan of *C. elegans* maintained on WT *E. coli* (Supplementary Figure 3). Together, these results suggest that folic acid shortens *C. elegans* lifespan on folate-deficient *E. coli* via AbgT-dependent uptake of PABA-glu during adulthood.

**Figure 4.**
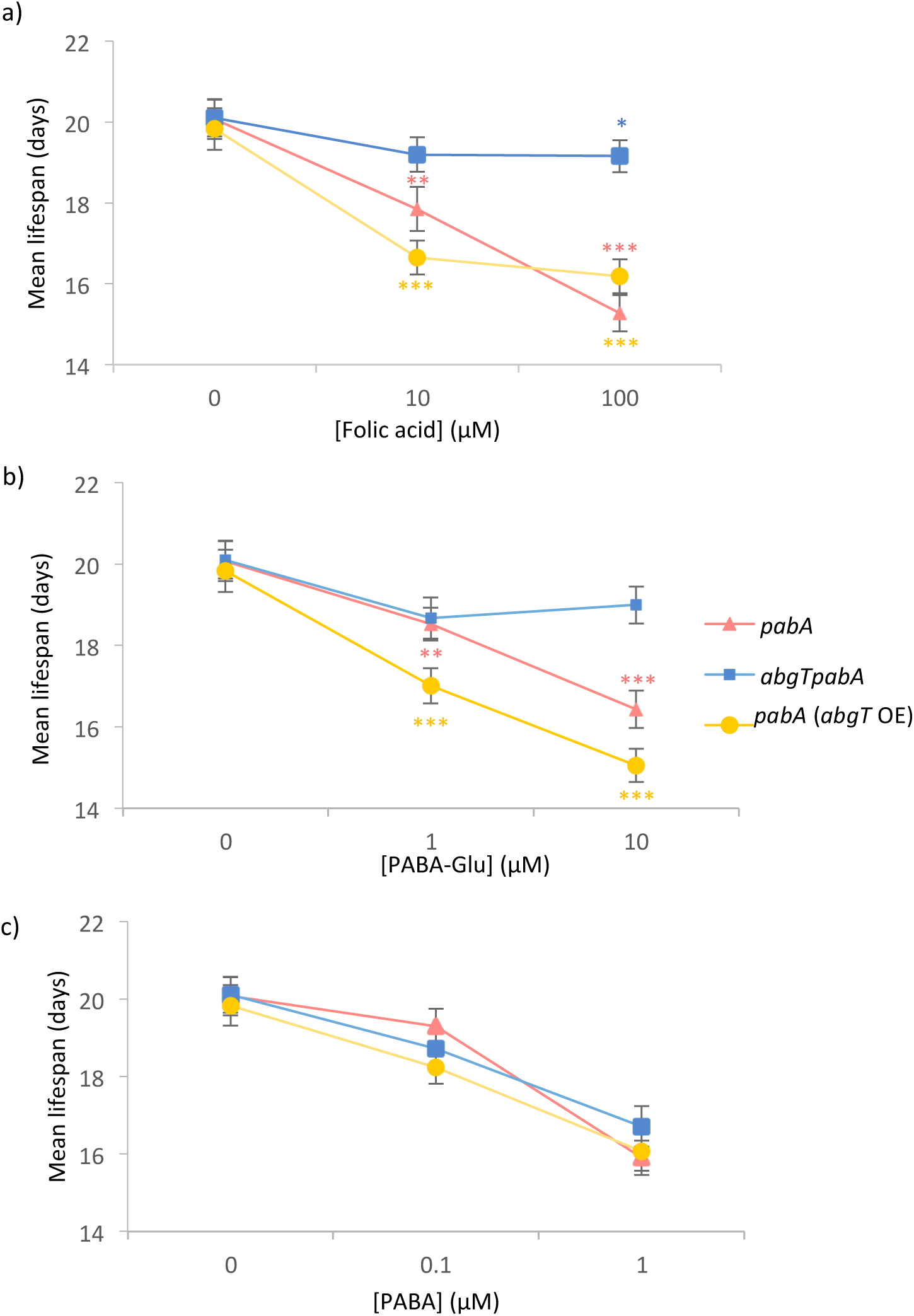
Folic acid shortens *C. elegans* lifespan via an *E. coli* abgT-dependent route during adulthood. Mean lifespan of WT *C. elegans* maintained from day 1 of adulthood on *pabA, abgTpabA,* or *pabA* (*abgT* OE) *E. coli* with supplementation of (a) folic acid (b) PABA-glu and (c) PABA. Error bars represent standard error. Asterisks denote the Log-rank non-parametric statistical test of survival, where: *P<0.05; **P<0.01; ***P<0.005 compared to lifespan on the non-supplemented condition of the same strain. Full lifespan data in Supplementary Table 1.

## DISCUSSION

In this study we have found that a major route of folic acid uptake by *C. elegans* is via *E. coli.* This route is dependent on the spontaneous breakdown of folic acid into PABA-glu and salvage of this product by the *E. coli* PABA-glu transporter, AbgT. We found that this route led to increases in the levels of several folate species in *E. coli* following folic acid supplementation. This route can also lead to folic acid shortening *C. elegans* lifespan when worms are cultured on bacteria with impaired folate synthesis. The increase in folate synthesis caused by folic acid supplementation leads to a bacterial activity/toxicity that is harmful to the worm over the long term (Figure 5).

**Figure 5.**
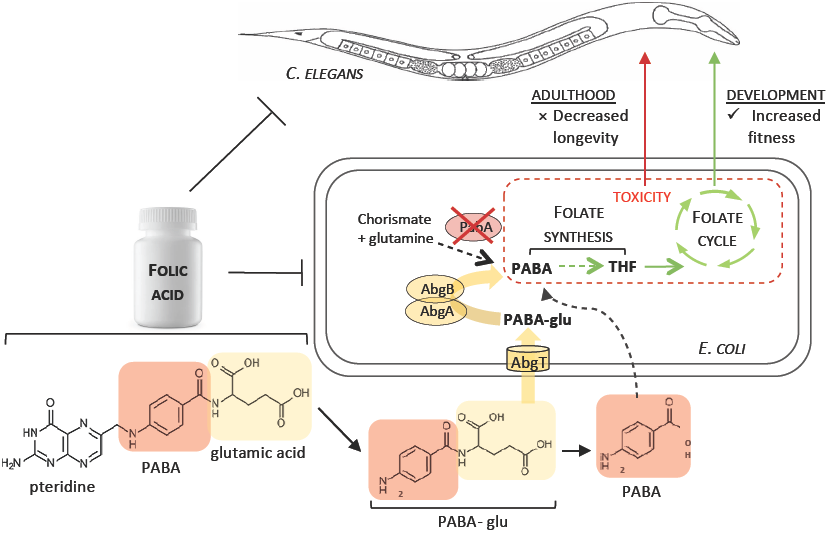
Schematic of the impact of folic acid supplementation on *C. elegans* via indirect uptake of breakdown products by *E. coli.* Folic acid is not taken up well by *C. elegans* directly. We find that the major uptake of folic acid by *C. elegans* is dependent on its breakdown into PABA-glu and uptake by the *E. coli* AbgT transporter. This route increases bacterial folate synthesis in both wild-type and *pabA* mutant *E. coli.* Under conditions of folate-deficiency *(pabA* mutant *E. coli),* increasing bacterial folate via this route is beneficial for *C. elegans* development. During *C. elegans* adulthood, this route has a negative impact on longevity as it promotes a bacterial folate-dependent toxicity.

Is it possible that this bacterial route for folic acid exists in humans? As far as we know, no other studies have tested folic acid supplements for the presence of PABA-glu or PABA, but several studies have reported issues with the stability and dissolution of commercial folic acid supplements (Hoag et al., 1997; Sculthorpe et al., 2001). In light of these issues, manufacturers have adopted a policy of ‘overages’ in order to ensure sufficient folic acid is released in the small intestine following ingestion (Andrews et al., 2017). The presence of PABA-glu and PABA in a commercially available folic acid source (Figure 3) combined the instability of folic acid at the low pH conditions of the stomach, makes it likely that PABA-glu and PABA will be available to the gut microbiota following supplementation. PABA has been identified as a human fecal excretory product after ingestion of folic acid (Denko et al.). The *abgT* gene is widespread in the human gut microbiota and studies in rodents (Rong et al., 1991) and piglets (Asrar and O'Connor, 2005) have demonstrated that infusion of labelled PABA into the cecum results in the incorporation of bacterially synthesized folate in host tissues. Thus the literature suggests that the components for such route exists in humans, but whether this accounts for significant amounts of folic acid is yet to be seen. We could only detect this route in *C. elegans* because of the poor bioavailability of folic acid in our folate deficiency model. Folic acid is taken up well by humans and leads to increases in serum folate levels, but folic acid is often found in the oxidised form, suggesting that it is not always bioavailable (Bailey et al., 2015; Gregory et al., 2005; Patanwala et al., 2014).

Further studies are required in order to determine whether folic acid supplementation affects the folate status of human gut microbes and whether this in turn impacts host health. Interestingly, there are several diseases associated with an increased abundance of folate-synthesizing gut bacteria, such as inflammatory bowel disease (Shin et al., 2015) and small intestinal bacterial overgrowth (Camilo et al., 1996), but a causal relationship between bacterial folate and disease has not been established. The abgT gene is found in the genomes of several enteric pathogens, including *Enterobacter cloacae, N. gonorrhoeae, Salmonella enterica, Shigella boydii* and *Staphylococcus aureus* in addition to *E. coli.* Whilst there is much consideration about the consequences of folic acid supplementation (Bailey et al., 2015; Boyles et al., 2016; Public_Health_England, 2017), our work here indicates that folic acid supplement instability and bacterial genotype are previously unexplored variables that may impact human health and thus warrant consideration in future reports and studies.

## MATERIALS AND METHODS

### Folates and related compounds

Folic acid, Folinic acid, PABA-Glu, 5-formylTHF-Glu_3_, 5-methylTHF-Glu_3_, methotrexate-Glu_6_ were obtained from Schircks, Switzerland. PABA, Vitamin B12 and folic acid were from Sigma Aldrich and folic acid supplement from Boots, UK.

### Culture conditions

Defined media (DM) was prepared as described (Virk et al., 2016), except that 10 nM B12 was added. B12, folic acid and antibiotics are added post-autoclaving for agar plates. DM for liquid culture is filter-sterilised. 0.1 μM PABA added to the liquid DM media used to seed the plates in order to maintain bacterial growth (apart the growth experiments in Figure 2). 30 μl of 3 ml fresh overnight LB culture is used to inoculate 5ml DM (15 ml Falcon tube). 25 μg/ml kanamycin (50 ug/ml ampicillin if necessary) added to both LB and DM pre-incubation. DM liquid cultures are incubated for 18 hr at 37°C, 220 RPM.

All strains derived from the Keio collection (Baba et al., 2006). See table. WT-Kan (Virk et al., 2016). *abgTpabA* double mutant was made using P1 transduction protocol as described in (Moore, 2011). The *abgT* over-expression plasmid (pJ128) (Carter et al., 2007) and empty vector (puc19) (Yanisch-Perron et al., 1985) were transformed into appropriate strains.

### *C. elegans* strains used

SS104 *glp-4(bn2),* UF208 (wild-type), and UF209 *gcp-2.1(ok1004)* (Virk et al., 2016).

### *E. coli* preparation and growth assay

Table of *E. coli* strains used in this study

| Strain | Genotype | Plasmid | Characteristics | Source |
| --- | --- | --- | --- | --- |
| BW25113/<br>pGreen 0029 | WT | pGreen 0029 | kan<sup>r</sup> | Virk et al 2016 |
| JW3323-1 | $\Delta pabA$ | n/a | kan<sup>r</sup> | Baba et al 2006 |
| JW5822-1 | $\Delta abgT$ | n/a | kan<sup>r</sup> | Baba et al 2006 |
| CMabgTpabA | $\Delta abgT \Delta pabA$ | n/a | kan<sup>r</sup> | This study |
| CM1 | WT | pUC19‡, pGreen 0029 | kan<sup>r</sup>, amp<sup>r</sup> | This study |
| CM2 | $\Delta pabA$ | pUC19 | kan<sup>r</sup>, amp<sup>r</sup> | This study |
| CM3 | $\Delta abgT \Delta pabA$ | pUC19 | kan<sup>r</sup>, amp<sup>r</sup> | This study |
| CM4 | WT (*abgT* OE) | pJ128‡,<br>pGreen 0029 | kan<sup>r</sup>, amp<sup>r</sup> | This study |
| CM5 | $\Delta pabA$ (*abgT* OE) | pJ128 | kan<sup>r</sup>, amp<sup>r</sup> | This study |
| CM6 | $\Delta abgT \Delta pabA$ (*abgT* OE) | pJ128 | kan<sup>r</sup>, amp<sup>r</sup> | This study |

*E. coli* was prepared as follows for all *E. coli* and *C. elegans* experiments. 30 μl of an overnight LB culture of *E. coli* was transferred into 5ml DM and incubated for 18 hr at 37 °C, 220 RPM. 100 μl the DM culture was seeded onto DM agar plates and incubated at 25°C for 4 days. *E. coli* was removed by pipetting 1 ml M9 onto the plate and a glass spreader to scrape off the bacterial lawn. The bacterial suspension was pipetted into a 1.5 ml Eppendorf and the volume was recorded (v). Tubes were vortexed vigorously to obtain a homogenised solution. 100 μl was taken and diluted with 900 μl M9 in a cuvette. A spectrophotometer was used to read bacterial growth at 600 nm. Bacterial growth was calculated by multiplying OD_600_ by the volume of the sample (v).

### *E. coli* folate extraction

Bacterial lawns were scraped from plates into micro centrifuge tubes using M9 solution and kept on ice. Volume (v), multiplied by the OD_600_ of the solution (diluted 1:5) gives a measure of the amount of material. Samples were concentrated in chilled microcentrifuge and pellets were snap frozen in liquid nitrogen. Pellets were thawed and resuspended in a volume of ice-cold 90% methanol: 10% folate extraction buffer (FEB: 50 mM HEPES, 50 mM CHES, 0.5% w/v ascorbic acid, 0.2 M DTT, pH 7.85 with NaOH) in proportion to bacterial content (37.5 × OD_600_ × v). FEB is spiked with 10 nM methotrexate-Glu_6_ as an internal standard. Samples were vortexed vigorously and left on ice for 15 min before centrifugation in a cooled microcentrifuge for 15 min at full speed. Supernatants were used for analysis.

### Folate LC-MS/MS analysis

Folates were detected by multiple reaction monitoring (MRM) analysis using a SCIEX QTRAP 6500 instrument. MRM conditions for folic acid, PABA, PABA-Glu, 5-Me-H_4_PteGlu_3_ (5-methylTHF-Glu_3_) and 5/10-CHO-H_4_PteGlu_3_ (formyl THF_3_) were optimised by infusion of standards into the instrument. The optimised conditions for –Glu_3_ folates were applied to other higher folates using MRM transitions described by Lu *et al.,* 2007 (Lu et al., 2007). Further confirmation of identity for folates of interest was achieved by performing enhanced product ion scans and comparing the fragment spectra with known standards.

The QTRAP 6500 was operated in ESI+ mode and was interfaced with a Shimadzu Nexera UHPLC system. Samples were separated using a Thermo PA2 C18 column (2.2 μm, 2.1 × 100 mm) with a gradient of 0.1% formic acid in water (mobile phase A) and acetonitrile (mobile phase B). Samples were maintained at 4°C and 2 μL aliquots were injected. The column was maintained at 40°C with a flow rate of 200 μL/min, starting at 2% B, held for 2 minutes, with a linear gradient to 100% B at 7 minutes, held for 1 minute, before a 7-minute re-equilibration step at 2% B that was necessary for consistent retention times. The column eluate flow to the MS was controlled via the QTRAP switching valve, allowing analysis between 4 and 8 minutes to minimise instrument contamination. Folates were quantified with reference to external standards and matrix effects were assessed by spiking of standards into extracted samples.

### Lifespan analysis

Survival analyses were performed as described (Virk et al., 2012). *glp-4(bn2)* worms were maintained at 15°C and shifted to 25°C at the L3 stage. At the L4/young adult stage, animals were placed on bacteria under the experimental conditions. All lifespan data is in Table S1. Statistical significance was determined using Log Rank and Wilcoxon tests of the Kaplan-Meier survival model.

## ACKNOWLEDGEMENTS

We thank the *C. elegans* Genetics Center, the *C. elegans* Knockout Consortium, and NBRP-*E.coli* at NIG for strains and we thank Sushmita Maitra and John Mathers for useful comments on the manuscript. This work was supported by a BBSRC DTP studentship.

## SUPPLEMENTARY INFORMATION

**Supplementary Figure 1.**
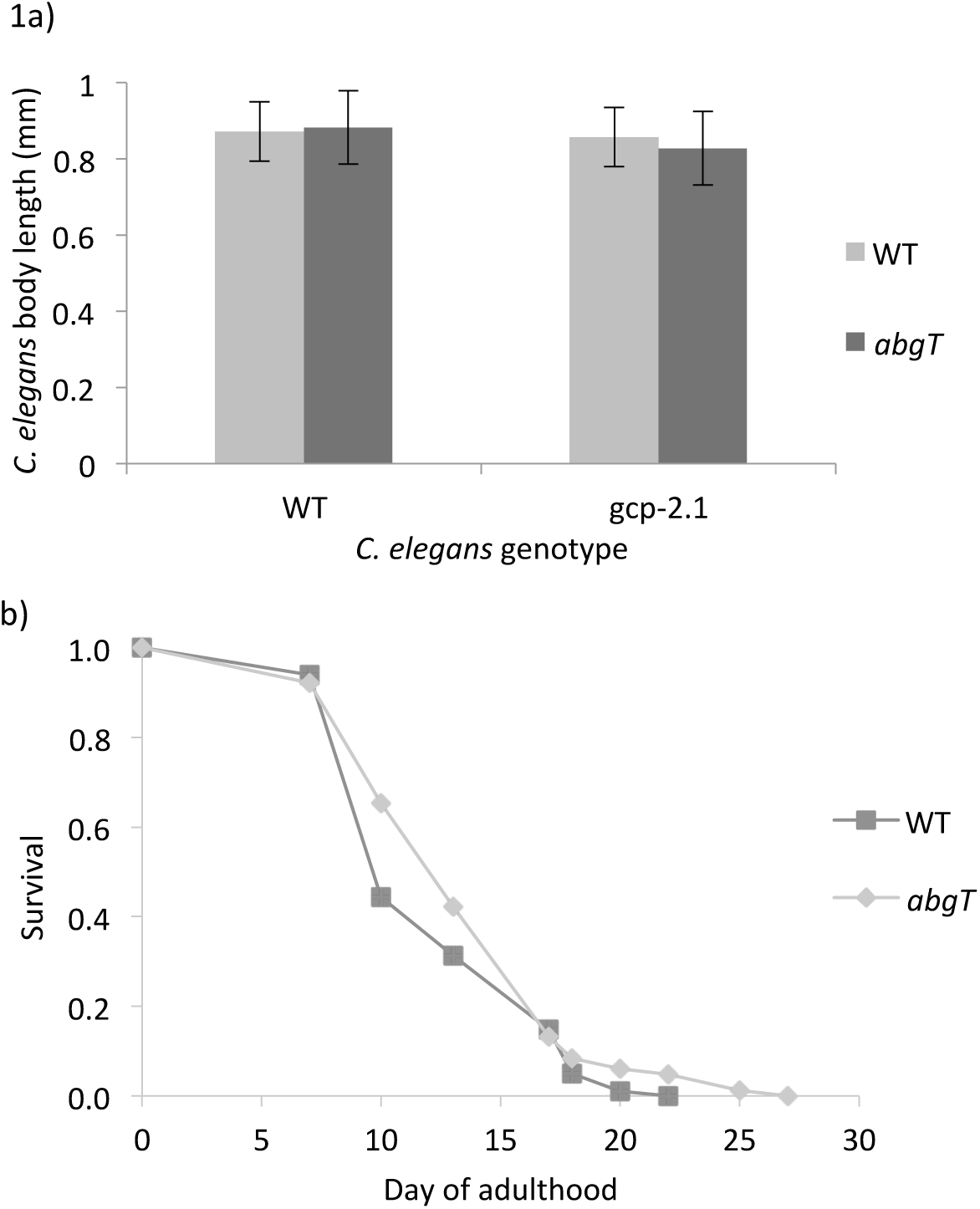
The *E. coli abgT* deletion has no effect on *C. elegans* development and lifespan. a) body length of wild-type and *gcp-2.1* mutant *C. elegans* at L4 stage raised on *abgT* mutant or wild-type *E. coli* (error bars represent standard deviation) b) survival curves of wild-type (*glp-4) C. elegans* on *E. coli* wild-type and *abgT* mutant. See Supplementary Table 1 for further details.

**Supplementary Figure 2.**
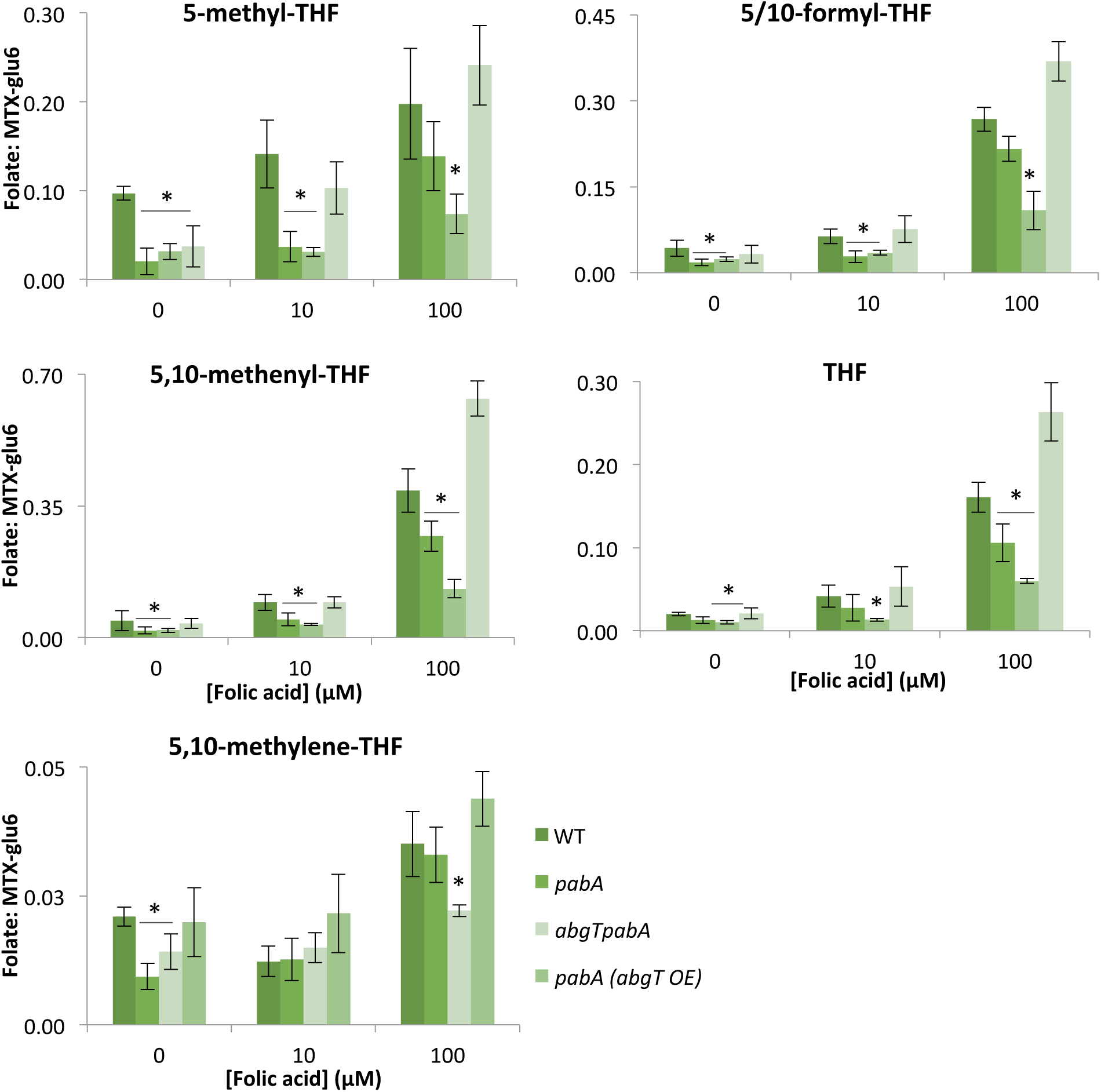
LC-MS/MS detects lower levels of THFs in *pabA* mutants, where *abgT* determines response to folic acid. Levels of THFs detected in extracts of WT, *pabA, abgTpabA, pabA (abgT* OE) mutants displayed as a ratio with an internal MTX-glu_6_ spike for normalization. Extracts were made after 4 days of bacterial growth at 25°C. Asterisks denote the test statistic from Student's *t* test comparison of means, where * = P<0.05. Error bars represent standard deviation over four replicates.

**Supplementary Figure 3.**
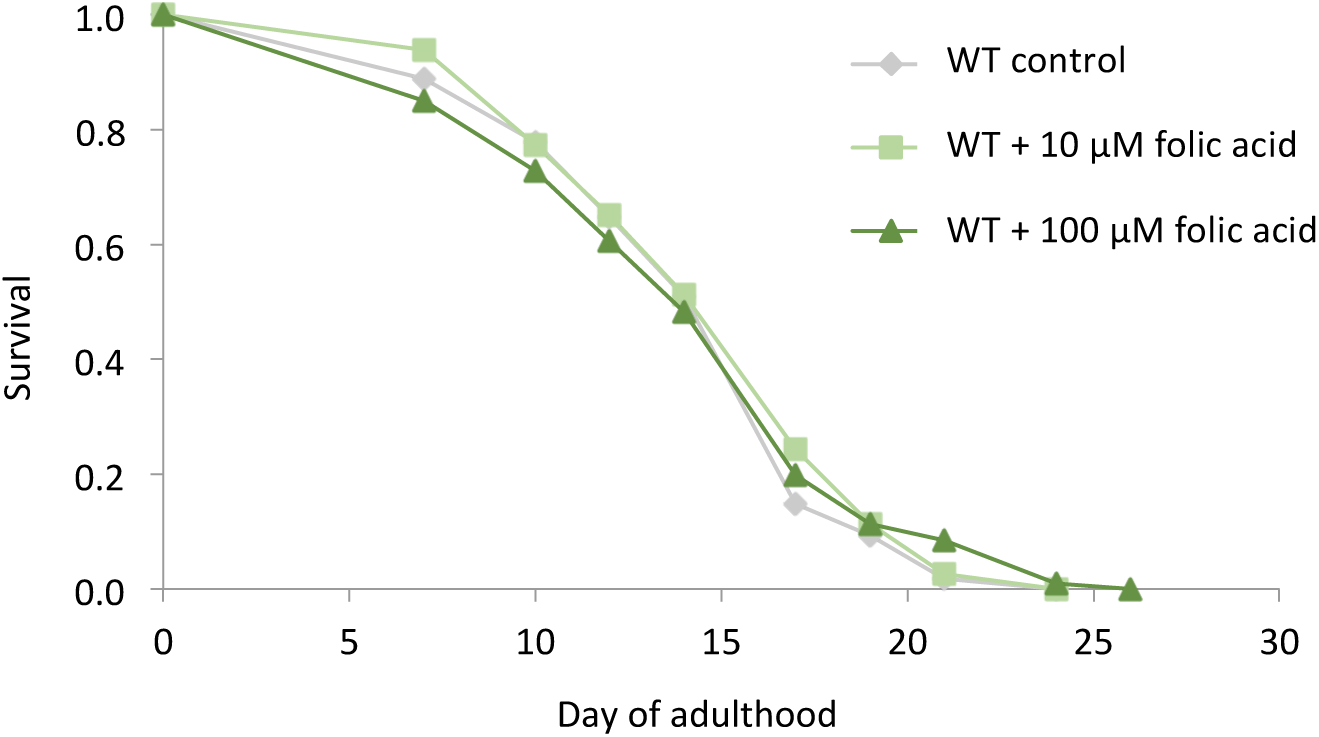
The impact of folic acid supplementation on *C. elegans* lifespan on wild-type *E. coli.* Survival curves of *C. elegans* maintained from day 1 of adulthood on plates supplemented with 10 μM and 100 μM folic acid.

**Supplementary Table 1:** **Summary of lifespan data** All lifespans conducted on defined media (DM) agar plates incubated at 25°C

| Fig. | Group name | n | Censor | Mean lifespan | Std. error | % change | p value (Log rank) | p value (Wilcoxon) | Supplement | Bacterial genotype | Plasmid | Antibiotics |
| --- | --- | --- | --- | --- | --- | --- | --- | --- | --- | --- | --- | --- |
| **5** | WT | 108 | 0 | 15.79 | 0.53 | n/a | n/a | n/a | Alkaline control | CM1 | pGreen 0029, pUC19 | 50 µg/ml carb, 25 µg/ ml kan |
|  | *pabA* | 94 | 3 | 20.08 | 0.52 | Control | n/a | n/a | Alkaline control | CM2 | pUC19 | 50 µg/ml carb, 25 µg/ ml kan |
|  | *pabA* + 0.1 µM PABA | 107 | 0 | 19.30 | 0.46 | -3.89% | 0.0973 | 0.3296 | 0.1 µM PABA | CM2 | pUC19 | 50 µg/ml carb, 25 µg/ ml kan |
|  | *pabA* + 1 µM PABA | 118 | 0 | 15.91 | 0.45 | -20.78% | <.0001* | <.0001* | 1 µM PABA | CM2 | pUC19 | 50 µg/ml carb, 25 µg/ ml kan |
|  | *pabA* + 1 µM PABA-Glu | 110 | 2 | 18.58 | 0.40 | -7.49% | 0.0030* | 0.0277* | 1 µM PABA-Glu | CM2 | pUC19 | 50 µg/ml carb, 25 µg/ ml kan |
|  | *pabA* + 10 µM PABA-Glu | 125 | 10 | 16.43 | 0.46 | -18.16% | <.0001* | <.0001* | 10 µM PABA-Glu | CM2 | pUC19 | 50 µg/ml carb, 25 µg/ ml kan |
|  | *pabA* + 10 µM folic acid | 93 | 5 | 18.20 | 0.55 | -9.36% | 0.0052* | 0.0061* | 10 µM Schircks folic acid | CM2 | pUC19 | 50 µg/ml carb, 25 µg/ ml kan |
|  | *pabA* + 100 µM folic acid | 112 | 1 | 15.28 | 0.45 | -23.93% | <.0001* | <.0001* | 100 µM Schircks folic acid | CM2 | pUC19 | 50 µg/ml carb, 25 µg/ ml kan |
|  | *abgT**pabA* | 91 | 3 | 20.10 | 0.44 | Control | n/a | n/a | Alkaline control | CM3 | pUC19 | 50 µg/ml carb, 25 µg/ ml kan |
|  | *abgT**pabA* + 0.1 µM PABA | 91 | 6 | 18.72 | 0.49 | -6.88% | 0.0631 | 0.0477* | 0.1 µM pABA | CM3 | pUC19 | 50 µg/ml carb, 25 µg/ ml kan |
|  | *abgT**pabA* + 1 µM PABA | 102 | 1 | 16.71 | 0.45 | -16.86% | <.0001* | <.0001* | 1 µM pABA | CM3 | pUC19 | 50 µg/ml carb, 25 µg/ ml kan |
|  | *abgT**pabA* + 1 PABA-Glu | 101 | 11 | 18.11 | 0.52 | -9.94% | 0.1733 | 0.0076* | 1 µM PABA-Glu | CM3 | pUC19 | 50 µg/ml carb, 25 µg/ ml kan |
|  | *abgT**pabA* + 10 µM PABA-Glu | 106 | 4 | 19.00 | 0.46 | -5.49% | 0.1817 | 0.0512 | 10 µM PABA-Glu | CM3 | pUC19 | 50 µg/ml carb, 25 µg/ ml kan |
|  | *abgT**pabA* + 10 µM folic acid | 110 | 1 | 19.20 | 0.43 | -4.49% | 0.1901 | 0.114 | 10 µM Schircks folic acid | CM3 | pUC19 | 50 µg/ml carb, 25 µg/ ml kan |
|  | *abgT**pabA* + 100 µM folic acid | 100 | 3 | 19.16 | 0.40 | -4.67% | 0.0476* | 0.0743 | 100 µM Schircks folic acid | CM3 | pUC19 | 50 µg/ml carb, 25 µg/ ml kan |
|  | *pabA* (*abgT* OE) | 62 | 10 | 19.90 | 0.52 | Contol | n/a | n/a | Alkaline control | CM5 | pJ128 | 50 µg/ml carb, 25 µg/ ml kan |
|  | *pabA* (*abgT* OE) + 0.1 µM PABA | 121 | 1 | 18.55 | 0.45 | -6.72% | 0.0049* | 0.0184* | 0.1 µM PABA | CM5 | pJ128 | 50 µg/ml carb, 25 µg/ ml kan |
|  | *pabA* (*abgT* OE) + 1 µM PABA | 104 | 2 | 16.07 | 0.50 | -19.19% | <.0001* | <.0001* | 1 µM PABA | CM5 | pJ128 | 50 µg/ml carb, 25 µg/ ml kan |
|  | *pabA* (*abgT* OE) + 1 µM PABA-Glu | 104 | 1 | 17.01 | 0.43 | -14.47% | <.0001* | <.0001* | 1 µM PABA-Glu | CM5 | pJ128 | 50 µg/ml carb, 25 µg/ ml kan |
|  | *pabA* (*abgT* OE) + 10 µM PABA-Glu | 117 | 0 | 15.05 | 0.41 | -24.33% | <.0001* | <.0001* | 10 µM PABA-Glu | CM5 | pJ128 | 50 µg/ml carb, 25 µg/ ml kan |
|  | *pabA* (*abgT* OE) + 10 µM folic acid | 119 | 0 | 16.65 | 0.42 | -16.30% | <.0001* | <.0001* | 10 µM Schircks folic acid | CM5 | pJ128 | 50 µg/ml carb, 25 µg/ ml kan |
|  | *pabA* (*abgT* OE) + 100 µM folic acid | 117 | 0 | 16.19 | 0.42 | -18.61% | <.0001* | <.0001* | 100 µM Schircks folic acid | CM5 | pJ128 | 50 µg/ml carb, 25 µg/ ml kan |
| **Sup. 1** | WT | 91 | 9 | 13.85 | 0.44 | Control | n/a | n/a | n/a | WT | pGreen 0029 | 25 µg/ml kan |
|  | *abgT* | 114 | 3 | 12.68 | 0.35 | -8.48% | 0.4312 | 0.0963 | n/a | JW5822-1 | n/a | 25 µg/ml kan |
| **Sup. 2** | WT | 108 | 0 | 14.58 | 0.40 | Control | n/a | n/a | Alkaline control | WT | pGreen 0029 | 25 µg/ ml kan |
|  | WT + 10 µM folic acid | 115 | 0 | 15.00 | 0.39 | 2.86% | 0.4448 | 0.4588 | 10 µM Schircks folic acid | WT | pGreen 0029 | 25 µg/ ml kan |
|  | WT + 100 µM folic acid | 106 | 1 | 14.57 | 0.48 | -0.10% | 0.566 | 0.844 | 100 µM Schircks folic acid | WT | pGreen 0029 | 25 µg/ ml kan |
| **Repeat expt.**<br>**(not shown)** | WT | 102 | 9 | 15.96 | 0.48 | Control | n/a | n/a | Alkaline control | CM1 | pGreen 0029, pUC19 | 50 µg/ml carb, 25 µg/ ml kan |
|  | WT + 10 µM folic acid | 98 | 12 | 15.96 | 0.58 | 0.03% | 0.8268 | 0.4283 | 10 µM Schircks folic acid | CM1 | pGreen 0029, pUC19 | 50 µg/ml carb, 25 µg/ ml kan |
|  | WT + 100 µM folic acid | 102 | 18 | 16.01 | 0.48 | 0.36% | 0.866 | 0.7792 | 100 µM Schircks folic acid | CM1 | pGreen 0029, pUC19 | 50 µg/ml carb, 25 µg/ ml kan |
|  | *pabA* | 113 | 10 | 18.35 | 0.56 | Control | n/a | n/a | Alkaline control | CM2 | pUC19 | 50 µg/ml carb, 25 µg/ ml kan |
|  | *pabA* + 10 µM folic acid | 90 | 11 | 17.44 | 0.64 | -4.95% | 0.2527 | 0.2605 | 10 µM Schircks folic acid | CM2 | pUC19 | 50 µg/ml carb, 25 µg/ ml kan |
|  | *pabA* + 100 µM folic acid | 122 | 3 | 15.86 | 0.40 | -13.57% | <.0001* | 0.0019* | 100 µM Schircks folic acid | CM2 | pUC19 | 50 µg/ml carb, 25 µg/ ml kan |
|  | *abgT**pabA* | 136 | 14 | 18.93 | 0.50 | Control | n/a | n/a | Alkaline control | CM3 | pUC19 | 50 µg/ml carb, 25 µg/ ml kan |
|  | *abgT**pabA* + 10 µM folic acid | 115 | 26 | 19.22 | 0.52 | 1.51% | 0.6416 | 0.5684 | 10 µM Schircks folic acid | CM3 | pUC19 | 50 µg/ml carb, 25 µg/ ml kan |
|  | *abgT**pabA* + 100 µM folic acid | 93 | 11 | 19.26 | 0.60 | 1.71% | 0.5868 | 0.4482 | 100 µM Schircks folic acid | CM3 | pUC19 | 50 µg/ml carb, 25 µg/ ml kan |
|  | *pabA* (*abgT* OE) | 101 | 23 | 18.42 | 0.56 | Control | n/a | n/a | Alkaline control | CM5 | pJ128 | 50 µg/ml carb, 25 µg/ ml kan |
|  | *pabA* (*abgT* OE) + 10 µM folic acid | 124 | 0 | 15.84 | 0.47 | -14.00% | 0.0005* | 0.0035* | 10 µM Schircks folic acid | CM5 | pJ128 | 50 µg/ml carb, 25 µg/ ml kan |
|  | *pabA* (*abgT* OE) + 100 µM folic acid | 105 | 0 | 15.77 | 0.50 | -14.36% | 0.0003* | 0.0041* | 100 µM Schircks folic acid | CM5 | pJ128 | 50 µg/ml carb, 25 µg/ ml kan |
|  | WT (*abgT* OE) | 110 | 9 | 16.11 | 0.50 | Control | n/a | n/a | Alkaline control | CM4 | pGreen 0029, pJ128 | 50 µg/ml carb, 25 µg/ ml kan |
|  | WT (*abgT* OE) + 10 µM folic acid | 99 | 4 | 16.05 | 0.57 | -0.37% | 0.9479 | 0.8905 | 10 µM Schircks folic acid | CM4 | pGreen 0029, pJ128 | 50 µg/ml carb, 25 µg/ ml kan |
|  | WT (*abgT* OE) + 100 µM folic acid | 127 | 4 | 15.78 | 0.39 | -2.02% | 0.3698 | 0.8168 | 100 µM Schircks folic acid | CM4 | pGreen 0029, pJ128 | 50 µg/ml carb, 25 µg/ ml kan |

## References

Andrews, K.W., Roseland, J.M., Gusev, P.A., Palachuvattil, J., Dang, P.T., Savarala, S., Han, F., Pehrsson, P.R., Douglass, L.W., Dwyer, J.T., et al. (2017). Analytical ingredient content and variability of adult multivitamin/mineral products: national estimates for the Dietary Supplement Ingredient Database. Am J Clin Nutr 105, 526–539, http://www.ncbi.nlm.nih.gov/pubmed/27974309

Asrar, F.M., and O’Connor, D.L. (2005). Bacterially synthesized folate and supplemental folic acid are absorbed across the large intestine of piglets. J Nutr Biochem 16, 587–593, http://www.ncbi.nlm.nih.gov/entrez/query.fcgi?cmd=Retrieve&db=PubMed&dopt=Citation&list_uids=16081276

Baba, T., Ara, T., Hasegawa, M., Takai, Y., Okumura, Y., Baba, M., Datsenko, K.A., Tomita, M., Wanner, B.L., and Mori, H. (2006). Construction of Escherichia coli K-12 in-frame, single-gene knockout mutants: the Keio collection. Mol Syst Biol 2, 2006 0008, http://www.ncbi.nlm.nih.gov/entrez/query.fcgi?cmd=Retrieve&db=PubMed&dopt=Citation&listuids=16738554

Bailey, L.B., Stover, P.J., McNulty, H., Fenech, M.F., Gregory, J.F., Mills, J.L., Pfeiffer, C.M., Fazili, Z., Zhang, M., Ueland, P.M., et al. (2015). Biomarkers of Nutrition for Development—Folate Review. The Journal of Nutrition 145, 1636S–1680S, http://jn.nutrition.org/content/145/7/1636S.abstract

Balamurugan, K., Ashokkumar, B., Moussaif, M., Sze, J.Y., and Said, H.M. (2007). Cloning and functional characterization of a folate transporter from the nematode Caenorhabditis elegans. Am J Physiol Cell Physiol 293, C670–681, http://www.ncbi.nlm.nih.gov/entrez/query.fcgi?cmd=Retrieve&db=PubMed&dopt=Citation&list_uids=17475669

Boyles, A.L., Yetley, E.A., Thayer, K.A., and Coates, P.M. (2016). Safe use of high intakes of folic acid: research challenges and paths forward. Nutrition Reviews 74, 469–474, http://dx.doi.org/10.1093/nutrit/nuw015

Camilo, E., Zimmerman, J., Mason, J.B., Golner, B., Russell, R., Selhub, J., and Rosenberg, I.H. (1996). Folate synthesized by bacteria in the human upper small intestine is assimilated by the host. Gastroenterology 110, 991–998, http://www.ncbi.nlm.nih.gov/pubmed/8613033

Carter, E.L., Jager, L., Gardner, L., Hall, C.C., Willis, S., and Green, J.M. (2007). Escherichia coli abg genes enable uptake and cleavage of the folate catabolite p-aminobenzoyl-glutamate. J Bacteriol 189, 3329–3334, http://www.ncbi.nlm.nih.gov/entrez/query.fcgi?cmd=Retrieve&db=PubMed&dopt=Citation&list_uids=17307853

Denko, C.W., Grundy, W.E., and et al. (1946). The excretion of B-complex vitamins by normal adults on a restricted intake.

Ducker, G.S., and Rabinowitz, J.D. (2017). One-Carbon Metabolism in Health and Disease. Cell Metabolism 25, 27–42, http://www.sciencedirect.com/science/article/pii/S1550413116304259

Green, J.M., and Matthews, R.G. (2007). Folate Biosynthesis, Reduction, and Polyglutamylation and the Interconversion of Folate Derivatives. EcoSal Plus 2, http://www.ncbi.nlm.nih.gov/pubmed/26443588

Gregory, J.F., Quinlivan, E.P., and Davis, S.R. (2005). Integrating the issues of folate bioavailability, intake and metabolism in the era of fortification. Trends in Food Science & Technology 16, 229–240, http://www.sciencedirect.com/science/article/pii/S0924224405000762

Halsted, C.H., Ling, E.H., Luthi-Carter, R., Villanueva, J.A., Gardner, J.M., and Coyle, J.T. (1998). Folylpoly-gamma-glutamate carboxypeptidase from pig jejunum. Molecular characterization and relation to glutamate carboxypeptidase II. J Biol Chem 273, 20417–20424, http://www.ncbi.nlm.nih.gov/pubmed/9685395

Han, B., Sivaramakrishnan, P., Lin, C.J., Neve, I.A.A., He, J., Tay, L.W.R., Sowa, J.N., Sizovs, A., Du, G., Wang, J., et al. (2017). Microbial Genetic Composition Tunes Host Longevity. Cell 169, 1249–1262 e1213, http://www.ncbi.nlm.nih.gov/pubmed/28622510

Hoag, S.W., Ramachandruni, H., and Shangraw, R.F. (1997). Failure of prescription prenatal vitamin products to meet USP standards for folic acid dissolution. J Am Pharm Assoc (Wash) NS37, 397–400, http://www.ncbi.nlm.nih.gov/pubmed/9519649

Kim, Y.-I. (2007). Folate and colorectal cancer: An evidence-based critical review. Molecular Nutrition & Food Research 51, 267–292, http://dx.doi.org/10.1002/mnfr.200600191

Lu, W., Kwon, Y.K., and Rabinowitz, J.D. (2007). Isotope ratio-based profiling of microbial folates. J Am Soc Mass Spectrom 18, 898–909, http://www.ncbi.nlm.nih.gov/entrez/query.fcgi?cmd=Retrieve&db=PubMed&dopt=Citation&list_uids=17360194

Marean, A., Graf, A., Zhang, Y., and Niswander, L. (2011). Folic acid supplementation can adversely affect murine neural tube closure and embryonic survival. Hum Mol Genet 20, 3678–3683, http://www.ncbi.nlm.nih.gov/pubmed/21693562

Nickerson, W.J., and Webb, M. (1956). Effect of folic acid analogues on growth and cell division of nonexacting microorganisms. J Bacteriol 71, 129–139, http://www.ncbi.nlm.nih.gov/pubmed/13295184

Patanwala, I., King, M.J., Barrett, D.A., Rose, J., Jackson, R., Hudson, M., Philo, M., Dainty, J.R., Wright, A.J., Finglas, P.M., et al. (2014). Folic acid handling by the human gut: implications for food fortification and supplementation. Am J Clin Nutr 100, 593–599, http://www.ncbi.nlm.nih.gov/pubmed/24944062

Pickell, L., Brown, K., Li, D., Wang, X.L., Deng, L., Wu, Q., Selhub, J., Luo, L., Jerome-Majewska, L., and Rozen, R. (2011). High intake of folic acid disrupts embryonic development in mice. Birth Defects Res A Clin Mol Teratol 91, 8–19, http://www.ncbi.nlm.nih.gov/pubmed/21254354

Public_Health_England (2017). Folic acid: updated SACN recommendations. https://www.gov.uk/government/publications/folic-acid-updated-sacn-recommendations

Rong, N., Selhub, J., Goldin, B.R., and Rosenberg, I.H. (1991). Bacterially synthesized folate in rat large intestine is incorporated into host tissue folyl polyglutamates. J Nutr 121, 1955–1959, http://www.ncbi.nlm.nih.gov/pubmed/1941259

Sculthorpe, N.F., Davies, B., Ashton, T., Allison, S., McGuire, D.N., and Malhi, J.S. (2001). Commercially available folic acid supplements and their compliance with the British Pharmacopoeia test for dissolution. J Public Health Med 23, 195–197, http://www.ncbi.nlm.nih.gov/pubmed/11585191

Shin, N.R., Whon, T.W., and Bae, J.W. (2015). Proteobacteria: microbial signature of dysbiosis in gut microbiota. Trends Biotechnol 33, 496–503, http://www.ncbi.nlm.nih.gov/pubmed/26210164

Smith, A.D., Kim, Y.-I., and Refsum, H. (2008). Is folic acid good for everyone? The American Journal of Clinical Nutrition 87, 517–533, http://ajcn.nutrition.org/content/87/3/517.abstract

Tran, P.V., and Nichols, B.P. (1991). Expression of Escherichia coli pabA. J Bacteriol 173, 3680–3687, http://www.ncbi.nlm.nih.gov/pubmed/2050628

Virk, B., Correia, G., Dixon, D.P., Feyst, I., Jia, J., Oberleitner, N., Briggs, Z., Hodge, E., Edwards, R., Ward, J., et al. (2012). Excessive folate synthesis limits lifespan in the C. elegans: E. coli aging model. BMC Biol 10, 67, https://doi.org/10.1186/1741-7007-10-67

Virk, B., Jia, J., Maynard, C.A., Raimundo, A., Lefebvre, J., Richards, S.A., Chetina, N., Liang, Y., Helliwell, N., Cipinska, M., et al. (2016). Folate Acts in E. coli to Accelerate C. elegans Aging Independently of Bacterial Biosynthesis. Cell Rep 14, 1611–1620, https://doi.org/10.1016/j.celrep.2016.01.051

Webb, M. (1955). Inactivation of analogues of folic acid by certain non-exacting bacteria. Biochim Biophys Acta 17, 212–225, http://www.ncbi.nlm.nih.gov/pubmed/13239661

Yanisch-Perron, C., Vieira, J., and Messing, J. (1985). Improved M13 phage cloning vectors and host strains: nucleotide sequences of the M13mp18 and pUC19 vectors. Gene 33, 103–119, http://www.ncbi.nlm.nih.gov/pubmed/2985470

